# A simplified, amplicon-based method for whole genome sequencing of human respiratory syncytial viruses

**DOI:** 10.1101/2022.11.24.516812

**Authors:** Xiaomin Dong, Yi-Mo Deng, Ammar Aziz, Paul Whitney, Julia Clark, Patrick Harris, Catherine Bautista, Anna Maria Costa, Gregory Waller, Andrew J Daley, Megan Wieringa, Tony Korman, Ian G. Barr

**Affiliations:** WHO Collaborating Centre for Reference and Research on Influenza, Royal Melbourne Hospital, at the Peter Doherty Institute for Infection and Immunity, Melbourne, VIC 3000, Australia; Department of Microbiology and Immunology, University of Melbourne, at the Peter Doherty Institute for Infection and Immunity, Melbourne, VIC 3000, Australia; Queensland Childrens Hospital, Brisbane, QLD 4101, Australia; Children’s Health Queensland Hospital & Health Service Brisbane, Brisbane, QLD 4101, Australia; UQ Centre for Clinical Research, Faculty of Medicine, University of Queensland, Herston, QLD 4029, Australia; Central Microbiology, Pathology Queensland, Royal Brisbane & Women’s Hospital, Herston QLD 4006, Australia; Department of Microbiology and Infectious Diseases, The Royal Children’s Hospital Melbourne, Parkville, VIC 3052, Australia; Department of Microbiology, Infection Prevention and Control, The Royal Children’s and Royal Women’s Hospitals, Parkville, VIC 3052, Australia; Monash Health, Clayton, VIC 3168, Australia

**Keywords:** respiratory syncytial virus, whole genome sequencing, one-step multiplex RT-PCR

## Abstract

Human Respiratory Syncytial Virus (RSV) infections pose a significant risk to human health worldwide, especially for young children. Whole genome sequencing (WGS) provides a useful tool for global surveillance to better understand the evolution and epidemiology of RSV and provide essential information that may impact on antibody treatments, antiviral drug sensitivity and vaccine effectiveness. Here we report the development of a rapid and simplified amplicon-based one-step multiplex reverse-transcription polymerase chain reaction (mRT-PCR) for WGS of both human RSV-A and RSV-B viruses. The method requires only two reactions for each sample, which significantly reduces the cost and time compared to other commonly used RSV WGS methods. In silico analysis and laboratory testing revealed that the primers used in the new method covered most of the currently circulating RSV-A and RSV-B. Amplicons generated were suitable for both Illumina and Oxford Nanopore Technologies (ONT) NGS platforms. This new method was tested on 200 clinical samples collected in Australia in 2020 and 2021 with RSV Ct values between 10 and 32. A success rate of 88% with a full coverage for the genome of 99 RSV-A and 77 RSV-B was achieved. This assay is simple to set up, robust, easily scalable in sample preparation and relatively inexpensive, and as such, provides a valuable addition to existing NGS RSV WGS methods.

## Introduction

Human Respiratory Syncytial Virus (RSV) belongs to the *Pneumoviridae* family and is a common cause of severe respiratory tract infections such as bronchiolitis and pneumonia in infants, young children, older adults and immunocompromised individuals ^1^. Despite the significant public health impact and global economic burden of RSV infections ^2, 3^, there are no vaccines or effective treatments currently available. The only prevention currently is an expensive monoclonal antibody prophylaxis for very young children ^4-6^.

RSV has a single stranded negative sense RNA genome of 15.2 kb in length comprising ten genes encoding transmembrane glycoproteins (F, G, SH), matrix proteins (M1, M2), proteins associated with the nucleocapsid (N, P, L) and two non-structural proteins (NS1, NS2) ^7^. RSV is divided into two major types RSV-A and RSV-B ^8^. Most genetic studies of RSV are focused on the attachment glycoprotein (*G*) gene, which is the most variable region and has been commonly used for RSV genotyping ^8^. In contrast, the F protein (which is responsible for fusion of viral and host cell membranes and syncytium formation) is highly conserved between strains and is the target of most vaccine and monoclonal antibody treatments currently being developed ^9, 10^. The mutations acquired during evolution may lead to changes in various aspects of virus biology, such as infectivity, antigenicity, drug resistance and potentially virulence ^11, 12^. Therefore, there is an ongoing need for effective monitoring of RSV viral evolution and spread throughout human society.

Next generation sequencing (NGS) technologies have been widely used for genomic studies of infectious diseases. Among different NGS approaches, the PCR amplicon-based NGS is a sensitive method for sequencing small genomes such as RSV for population-scale viral surveillance ^13^. One challenge associated with this method is the generation of PCR amplicons that can cover all variants. Several singleplex reverse-transcription (RT)-PCR amplicon-based NGS methods have been used for RSV whole-genome sequencing (WGS) including a commonly used method that requires at least four separate reactions per genome ^14, 15^. While the use of multiplex PCR (mPCR) to amplify multiple DNA fragments in the same reaction provides a simpler solution to reducing the number of reactions^16^, it often leads to a decreased sensitivity, due to complex primer-primer interactions and competition for usage of reagents ^17^. Therefore, the establishment of optimal primer pools for multiplex PCR can be challenging and may require many sets of primers as seen with the ARTIC protocol for COVID-19 WGS ^18, 19^. The development of amplicon-based WGS should ideally be suitable for multiple NGS platforms, especially in developing countries where the burden of RSV is greatest. One such system that is currently widely used is the ONT’s MinION system, which is a portable and low-cost NGS that can significantly simplify the sequencing workflow and shorten the turnaround time for sequencing ^20^.

In April 2019, the World Health Organization (WHO) initiated phase 2 of their Global RSV Surveillance program to study the circulation and impact of RSV worldwide. One of the elements of this study was to increase the sequencing capacity for RSV viruses ^1^, especially in developing countries. In order to meet the increasing demand for RSV sequencing, we endeavoured to develop a simplified one-step multiplex RT-PCR (mRT-PCR) method for RSV WGS. This approach uses a pool of primers to generate a number of overlapping PCR amplicons in two, one-step RT-PCR reaction tubes that is suitable for both RSV-A and RSV-B positive samples without knowing the RSV type. This method significantly simplifies the costly and labour-intensive steps associated with other currently used amplicon-based methods that use at least four separate reactions. This new method can be used to accelerate and streamline the sequencing of RSV genomes using a range of NGS platforms, including Illumina and ONT platforms.

## Results

### Development of one-step multiplex RT-PCR (mRT-PCR) for RSV WGS

Primers for mRT-PCR were designed based on the alignment of 1520 RSV-A and 1364 RSV-B genome sequences with nucleotides more than 14900 bp (submitted to GISAID from 1^st^ January 2015 to 9th July 2021 (Supplementary Fig. 1)). Conserved regions of both RSV-A and RSV-B were used for primer selection to allow the same primers to be used for both RSV-A and RSV-B amplification. The specificity of these primers was further checked by BLAST to ensure that the primers were specific for human RSV. Six pairs of primers were designed to amplify 6 amplicons with distinguishable sizes in two multiplexed PCR reactions for RSV whole genomes. Degenerate bases were introduced in the primers to cover some of the RSV variants. (Table 1). These primers were tested using a SuperScript IV (SSIV) One-Step RT-+PCR kit (Thermo Fisher Scientific) in two reactions on both RSV-A and RSV-B positive control samples. Primer concentrations were optimized so that roughly equal densities of PCR product could be amplified for each PCR primer pair. As shown in Fig. 1A, six amplicons in two mRT-PCR tubes that fully covered the whole RSV genome were amplified from both RSV-A (hRSV/A/Australia/QLD-RBWH114/2021) and RSV-B (hRSV/B/Australia/QLD-RBWH289/2021) respiratory clinical samples with relatively similar efficiency and the six amplicons were easily distinguishable by size using electrophoresis (Figure 1B). The amplicons from two reactions were combined in an equal amount and used for library preparation for iSeq100 NGS. Results revealed that both RSV-A and RSV-B genomes were fully sequenced with relatively even coverage (Figure 1C). The successful mRT-PCR results demonstrated that the primer combination worked well in the two reactions with no adverse interference with each other, suggesting that they could be used for sequencing both RSV-A and RSV-B.

**Table 1:**
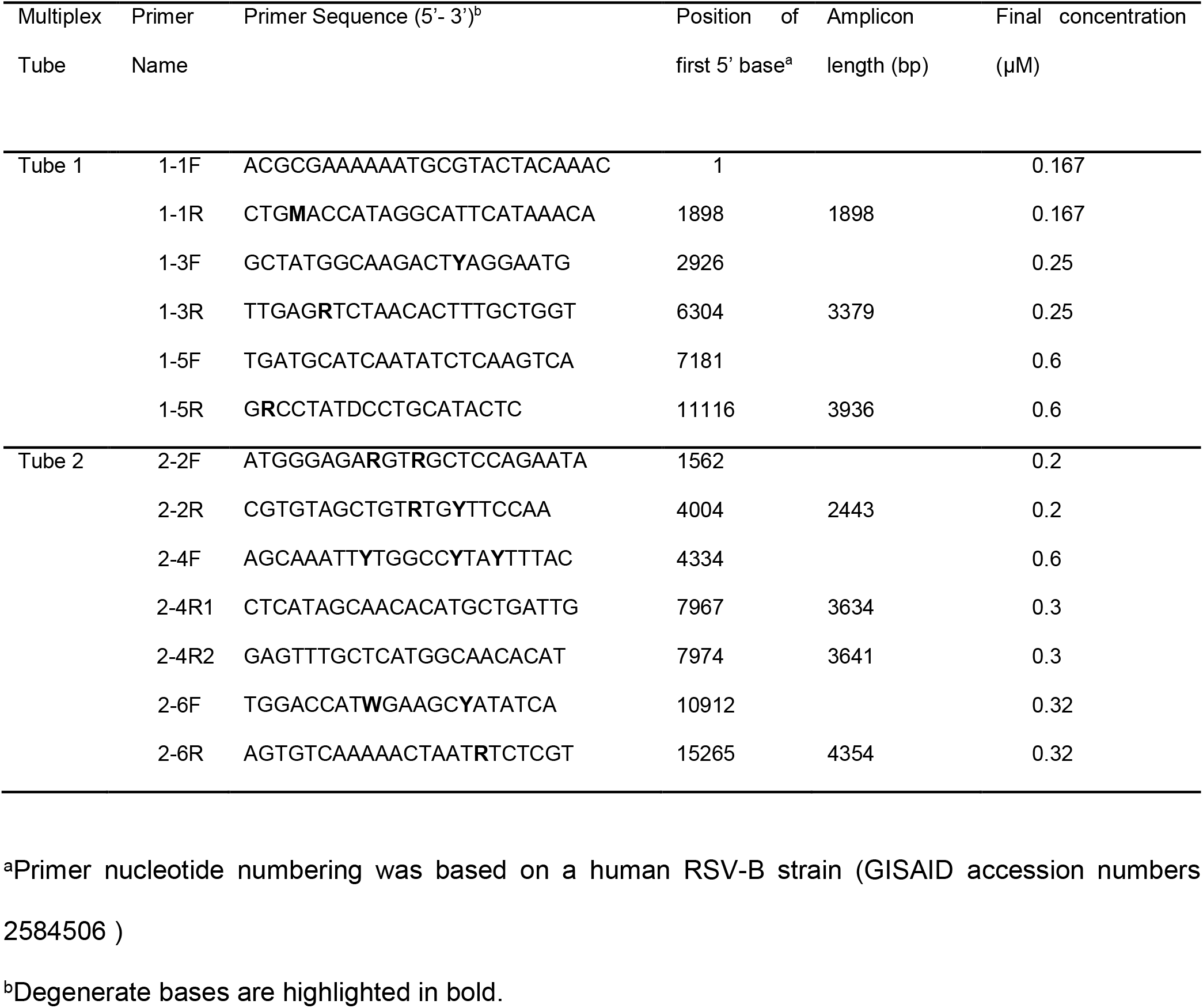
The information for primers used in the one-step multiplex RT-PCR

**Figure 1.**
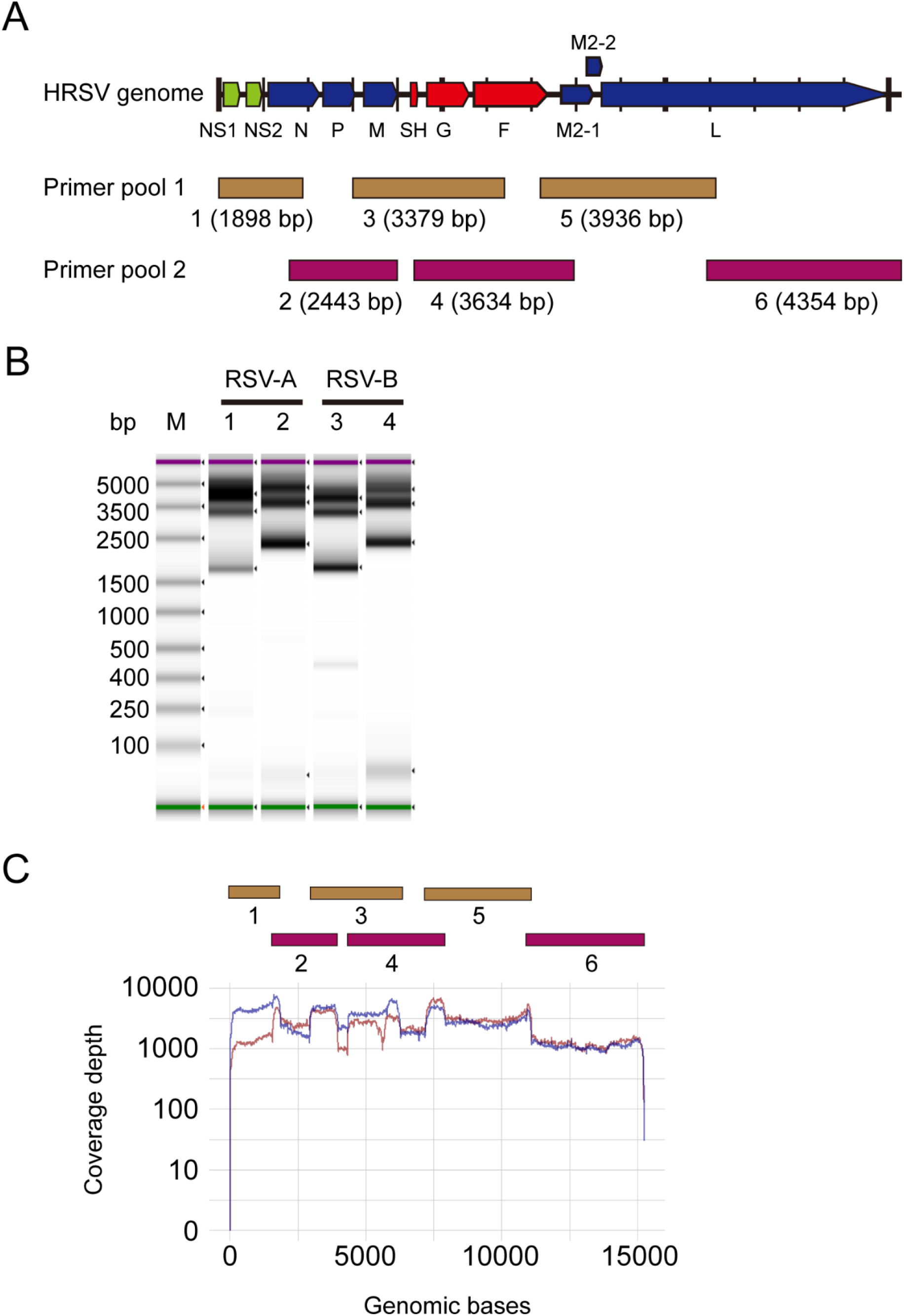
One-step multiplex RT-PCR for RSV WGS (A) Schematic overview of the 6 amplicons designed to cover the RSV genome and the encoded proteins; NS1: non-structure protein 1, NS2: non-structure protein 2, N: nucleoprotein, P: phosphoprotein, M: matrix protein, SH: small hydrophobic protein, G: glycoprotein, F: fusion protein, M2-1: Matrix protein 2-1, M2-2: Matrix protein 2-2, L: large polymerase. One PCR reaction contains pool 1 primers that generate three fragments (numbered as 1, 3 and 5) ranging from about 1898 to 3936 bp, while the second PCR reaction has pool 2 primers that generate another three fragments (numbered as 2, 4 and 6) of sizes between 2443 and 4354 bp based on a human RSV-B strain (GISAID accession numbers 2584506). (B) Representative Tapestation images of two-pool PCR products of viral RNA from one RSV-A and one RSV-B positive clinical samples. Molecular marker (M); mRT-PCR products of pool 1 (lane 1) and pool 2 (lane 2) reactions from hRSV/A/Australia/QLD-RBWH114/2021; mRT-PCR products of pool 1 (lane 3) and pool 2 (lane 4) from hRSV/B/Australia/QLD-RBWH289/2021. (C) Coverage depth of sequenced RSV-A (Dark red) and RSV-B (Dark blue) in genomic positions covered by six overlapping amplicons.

### Performance of RSV mRT-PCR in clinical samples

De-identified RSV positive respiratory clinical samples obtained from Australian laboratories collected between December 2020 and April 2021 were re-screened using the CDC RSV duplex assay^21^ and their Ct values were determined. Two hundred and sixteen RSV positive respiratory clinical samples with cycle thresholds (Ct) values between 10 and 32 were selected for RSV mRT-PCR testing; 50 RSV-A and 55 RSV-B samples with Ct values below 20 (105/115= 91.3%) had all six bands clearly detectable and were selected for WGS. As expected, all these samples had RSV whole genomes fully sequenced.

Next, we assessed the sensitivity of this new mRT-PCR assay for sequencing RSV clinical samples with relatively higher Ct values than those tested above. The mRT-PCR amplicons from 101 RSV samples with Ct values ranging from 20 to 32 were used for Illumina NGS, regardless of whether all six amplicons were visible or not upon TapeStation analysis. WGS was obtained from 57 out of 101 samples with Ct value up to 27 (56.4%) (Fig. 2), of these, all six PCR amplicons were only obvious in 27 samples, with at least one amplicon not detectable by TapeStation from the other 30 samples. This result suggested that NGS can still work well even with weak PCR fragments that are not visible on a TapeStation System. In addition, 24 samples generated partial RSV genomes, importantly 18 with complete *G* and *F* genes (GF) and 6 with the complete *G* gene only.

**Figure 2.**
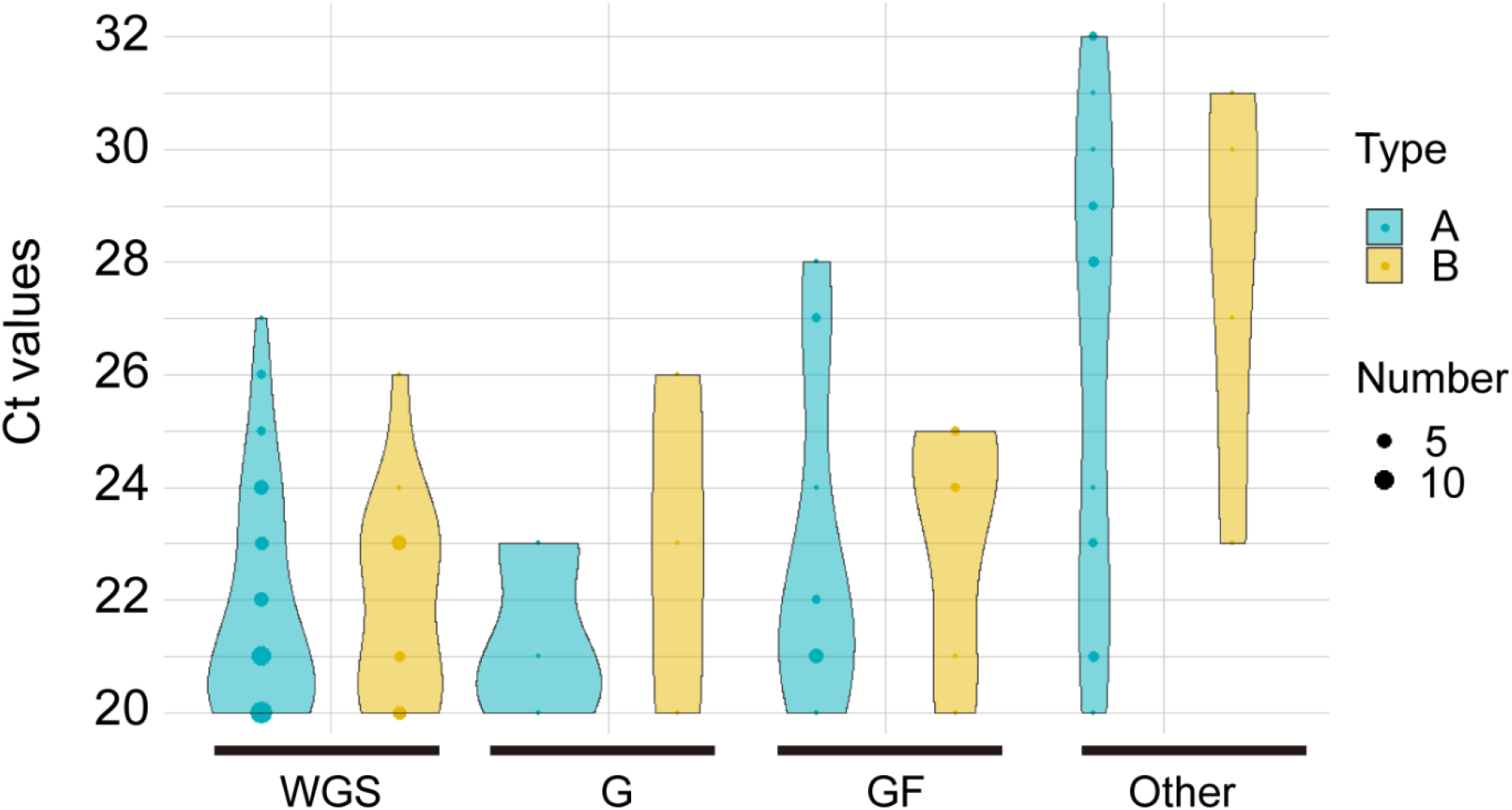
Sensitivity of RSV genome sequencing in clinical samples with varied Ct values Dot and violin plot showing the outcomes of RSV-A and RSV-B samples sequenced by NGS according to their RT-PCR Ct values; whole genome sequence obtained (WGS), *G* gene only obtained (G), *G* and *F* genes obtained (GF), and partial sequence without *G* gene obtained (Other).

### DNase treatment of samples with lower viral load improved NGS results

As shown in Supplementary Fig. 2A, non-specific amplicons were occasionally observed in the mRT-PCR products of some samples which resulted in NGS reads that were not from RSV, for examples, only 70.25 % (hRSV/A/Australia/QLD-RBWH008/2021) and 74.22 % (hRSV/B/Australia/QLD-RBWH150/2021) sequencing reads aligned to RSV-A and RSV-B reference sequences according to the iterative refinement meta-assembler (IRMA) analysis ^22^, respectively. To estimate read abundances at the species level, Kraken 2 ^23^, a taxonomic classification system, was used and revealed about 30% and 26% sequencing reads from these two samples respectively were human DNA sequences (Supplementary Fig. 2B and 2C), indicating the presence of human genome DNA in the extracted viral RNA. Moreover, the percentage of RSV specific reads decreased as the Ct value increased for all the above sequenced samples, according to a linear regression line plotted with polynomial interpolation (Supplementary Fig. 2D), suggesting that the samples with low RSV viral abundance were more significantly affected by human genome contamination than those with high RSV viral loads in this assay.

To improve sensitivity of the mRT-PCR NGS in samples with lower viral loads, we used DNase treatment on the purified viral RNA in samples with a RSV Ct value of 23 and above, and compared mRT-PCR and NGS results on these samples with and without DNase treatment. Thirty-one RSV-A and 17 RSV-B positive clinical samples were chosen for the DNase treatment comparison. Notably, DNase treatment of viral RNA significantly improved the efficiency of mRT-PCR, and RSV amplicons were more prominent upon TapeStation examination (Fig. 3, Supplementary Fig. 3A). Furthermore, WGS were obtained from 60.4% of the samples (29 samples) including 18 RSV-A and 11 RSV-B, while GF sequences were obtained from a further 12 viruses (25%) and one sample had the complete *G* gene only. In total, 42 samples with Ct values of up to 32 generated useful sequence data, amounting to 87.5% of the samples tested (Supplementary Fig. 3B), with only 6 samples (12.5%) failing to generate useful sequences with complete *G* gene. In contrast, without DNase treatment, WGS sequences could only be successfully generated from 21 samples (43.8%) including 14 RSV-A and 7 RSV-B, plus 8 samples with GF sequences (16.7%) and 3 with only G sequences (6.3%) generated amounting to a total success rate of 66.7% (Supplementary Fig. 3C) while 16 (33.3%) that did not produce useful sequence data. Hence, the DNase treatment improved the success rate of useful sequence generation by 20.8%.

**Figure 3.**
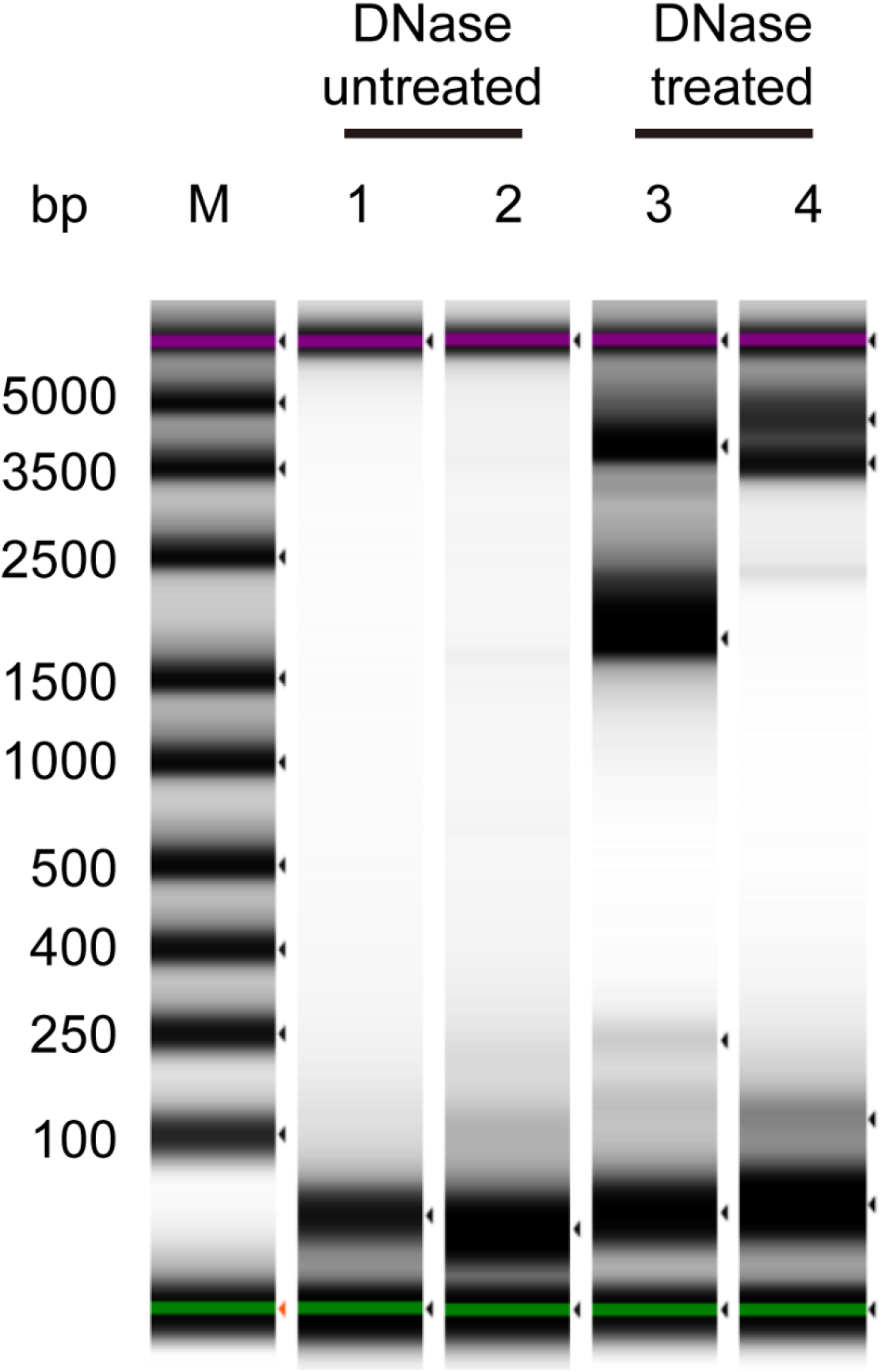
DNase treatment improved assay sensitivity RSV amplicons (pool 1 and 2) generated from a RSV-B positive clinical sample hRSV/B/Australia/QLD-RBWH239/2021 without and with DNase treatment.

NGS data analysis of all 48 samples indicated that the mean percentage of RSV specific reads from all samples was significantly increased from 25.6% to 88.9% after DNA removal (Supplementary Fig. 3D), whereas human reads abundance dropped from 60.9% to 5.7% as estimated by Kraken 2 (Supplementary Fig. 3E).

### Sequencing of samples on both Illumina and ONT platforms

We further evaluated sequencing of the mRT-PCR products on the Oxford Nanopore (ONT) MinION system. The same mRT-PCR products from two RSV positive clinical samples with high RSV viral loads (hRSV/A/Australia/QLD-RBWH060/2021; hRSV/B/Australia/QLD-RBWH289/2021) were used for sequencing on both Illumina and ONT platforms comparison. For long-reads sequencing with ONT, a ligation-based NGS library preparation method was used and a bead clean-up of PCR products step was added before preparing libraries to remove small fragments (<500 nts). The resulting datasets were analyzed for coverage plots and whole genome consensus sequences. As shown in Fig. 4A and 4B, both Illumina iSeq100 and ONT MinION generated data with good coverage throughout the whole genome. In comparison to Illumina iSeq sequencing, a relative higher depth across the entire genome region was achieved with ONT when sequenced for three hours (Fig. 4A and 4B). Moreover, a flat depth curve was observed within the same PCR amplicon region for ONT sequencing, indicating even coverage depth was achieved by ONT sequencing in these regions as its library preparation does not require fragmentation, which reduces any bias introduced during library preparation.

**Figure 4.**
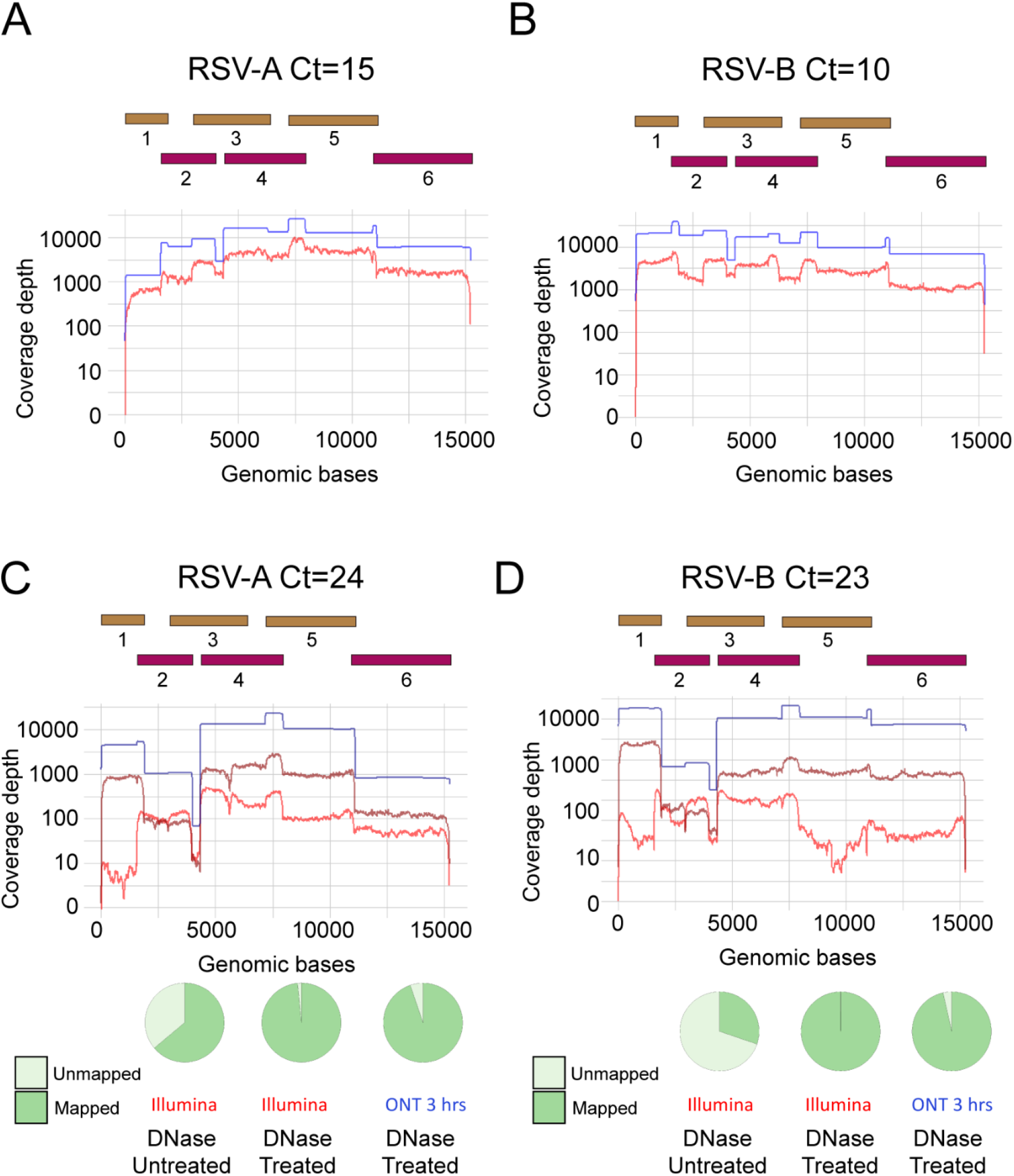
Coverage depths comparison of NGS data generated from Illumina and ONT platforms High coverage depths of a RSV-A (A) and a RSV-B (B) samples with high viral loads (low Ct values) sequenced on Illumina (red) and ONT (blue) platforms at each nucleotide position. A RSV-A (C) and a RSV-B (D) sample with low viral loads (high Ct values) were treated or not treated with DNase and run on the two platforms. Illumina sequencing without DNase treatment shown in red and after treatment shown in dark red. ONT sequencing after DNase shown in dark blue. The pie charts show the NGS reads that were RSV specific (green) and nonspecific (light green) from both iSeq100 and MinION platforms with or without DNase treatment. No data was obtained for samples of low viral loads without DNase treatment due to failure in generating enough amount of NGS library for ONT sequencing.

We further evaluated if DNase treatment could improve sequencing quality with ONT platform and compared the results obtained with the Illumina system. Both DNase treated and untreated viral RNA from two clinical samples with relatively low RSV viral loads (hRSV/A/Australia/QLD-RBWH191/2021 with a Ct of 24 and hRSV/B/Australia/QLD-RBWH239/2021 with a Ct of 23) were used for amplicon generation comparison with the new mRT-PCR assay (Fig. 3 and supplementary Fig. 3A). We found DNase treatment also improved the NGS performance on ONT platform as well as the Illumina platform (Fig. 4C and 4D). Notably, without DNase treatment, two samples failed to generate enough amount of NGS library for ONT sequencing, even though more PCR product was used for ONT library preparation than for the Illumina sequencing. This suggested that the NGS sensitivity of Illumina sequencing with on-bead tagmentation library preparation is higher than that of ONT sequencing with ligation-based library preparation in this study. The consensus sequences generated from Illumina iSeq and ONT MinION platforms were compared for accuracy. Sequence variations from 3 to 19 nucleotides were identified from the 4 samples from a total of 15kb sequence (Supplementary Table 1). A close examination revealed that the mismatches were mainly located in homopolymeric regions or in regions with a low sequencing depth. Importantly no mismatches were found in the *G* or *F* genes, which are the most important genes for typing, surveillance and vaccine/monoclonal antibody monitoring.

## Discussion

To the best of our knowledge, previously published amplicon-based RSV WGS approaches require at least 4 different singleplex PCR amplification reactions following the cDNA synthesis from viral RNA with separate reverse transcription reactions that takes significant time to process a large number of samples ^8, 14, 15^. The mRT-PCR method described herein provides a convenient, cost-effective and more scalable way for sequencing the RSV A or B whole genomes. It only requires two tubes and a one-step RT-PCR reaction to generate overlapping PCR fragments from viral RNA and therefore significantly reduces handling time and reduces the risk of contamination. Traditional multiplex PCR-tiling amplicon methods for WGS normally have primers designed to generate several amplicons of similar size for achieving comparable PCR amplification efficiencies in the same tube, making it impossible to tell if all amplicons have been successfully amplified without sequencing them. However, this mRT-PCR method generated amplicons are of easily distinguishable sizes, making the identification of all segments amplified very easy through electrophoresis, giving the user a good indication whether WGS can be likely achieved for each sample.

By introducing an optional DNase treatment step for original specimens with lower viral loads (and higher Ct values), we were able to improve the WGS success rate. Even if the WGS could not be obtained from some samples, the *G* gene sequence was obtained in many of these samples along with the *F* gene in some. The *G* and *F* genes are the most informative genes with the *G* gene being the more variable and being used for both evolutionary analysis as well as for classification systems that are important for RSV surveillance, while the F protein is the main target for vaccine development and for prophylactic monoclonal antibody preventatives and hence requires close scrutiny. Other studies have also demonstrated advantages in using DNase treatment for enhancing virus detection in serum and plasma samples by boosting the specific fragment generation with the sequence-independent single primer amplification (SISPA) for sequencing ^24^. Recently, DNase treatment on RNA extracted from clinical samples also demonstrated improved RSV detection with SISPA followed by NGS ^25^.

The method described here can be used to prepare libraries for different sequencing-platforms such as Illminia or ONT. According to a recent study of ONT sequencing of SARS-CoV-2, long amplicons (∼2 and ∼2.5-kb) PCR-tiling clearly showed better performance in coverage variation and overall quality of the final sequence consensus when compared to short amplicons (∼400-bp) ^26^. Thus, our method with long amplicons for RSV WGS is also suitable for ONT in addition to Illumina sequencing.

A limitation of this study is that although we have demonstrated that tiling amplicons generated by this approach can be sequenced either on the Illumina iSeq platform or on the long-read ONT MinION platform, we characterized this new method mostly using an Illumina platform and only several RSV whole genomes were sequenced on the ONT platform. An advantage of using the Illumina DNA preparation kits is that saturation-based DNA normalization is performed when doing on-bead tagmentation ^27^. Therefore, up to 48 NGS libraries were pooled for a multiplex sequencing on an Illumina iSeq device in this study, without individual library quantitation and normalization. However, for ONT sequencing, a ligation-based NGS library preparation method was used for long-read sequencing and it is highly recommended to do individual library quantitation and normalization before pooling them for sequencing, as we did in this study.

Overall, we have demonstrated a new method and workflow for NGS of RSV whole genomes directly from clinical respiratory samples without the need for typing that is based on a one-step mRT-PCR amplicon based approach (Fig. 5). For optimum NGS results, we suggest the pre-treatment of RNA with DNase prior to mRT-PCR for samples with RSV Ct values between 23 and 28. Downstream NGS could be done with either the short-read Illumina platforms or the long-read ONT sequencing platform. With its robust performance, faster and more scalable preparation, this protocol is a valuable addition to existing RSV WGS methods.

**Figure 5.**
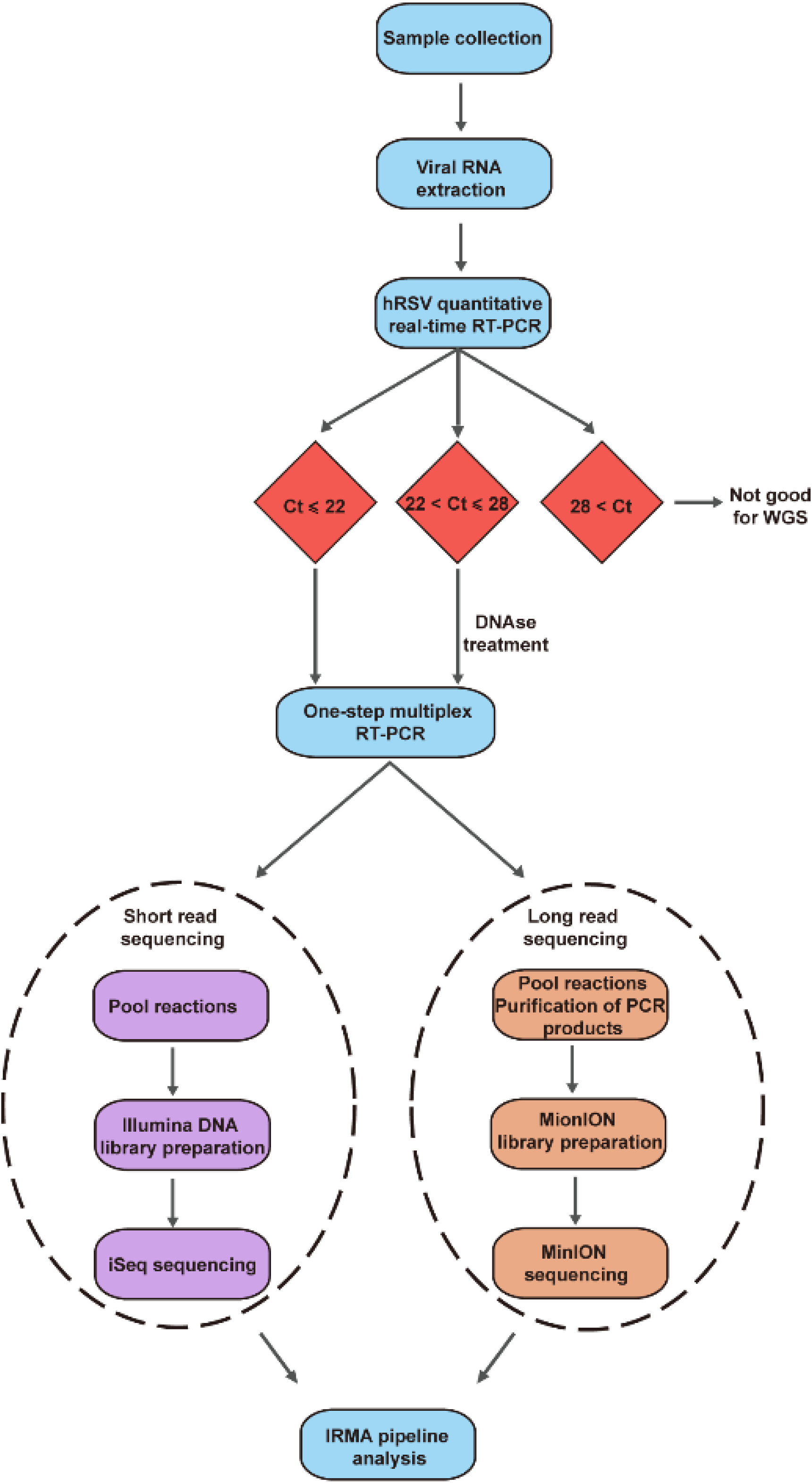
Schematic diagram of the RSV whole genome sequencing workflow A decision tree showing the different processing paths depending on the viral load in the clinical samples

## Methods and materials

### Clinical specimens and isolates

Samples used in this study were de-identified respiratory samples (including nasal swabs, nasopharyngeal swabs, nasal washes or nasopharyngeal aspirates) that were positive for RSV by routine clinical laboratory molecular testing at Queensland and Victorian (Australia) laboratories collected between December 2020 and April 2021. An aliquot of these samples was shipped to the Centre as part of the WHO RSV surveillance program and samples were stored at -80°C until further analyzed.

### RNA extraction and real-time RT-PCR assay

Viral RNA was extracted from 200 μl of clinical sample using QIAamp 96 Virus QIAcube HT (QIAGEN) according to the manufacturer’s instruction. The CDC Respiratory Syncytial Virus Real-time RT-PCR panel (RSV_RUO-01) ^21^, was used for RSV subtype detection with TaqPath 1-Step Multiplex Master Mix (No ROX) on Applied Biosystems 7500 Real-Time PCR System (ABI, Life Technologies, Waltham, MA) as instructed. Reactions were run using the following cycling conditions: one cycle of 25°C for 2 min, followed by one cycle of 53°C for 10 min and 95°C for 2 min, followed by 40 cycles of 95°C for 3 s and 60°C for 1 min. The resulting cycle threshold (Ct) values were rounded to the nearest whole number.

### RSV amplicon generation by one-step multiplex RT-PCR for RSV WGS

SuperScript IV One-Step RT-PCR System (Invitrogen) was used for RSV whole genome amplification with two separate multiplex RT-PCR reactions using six pairs of primers designed for this study. Briefly, 50 µL reaction containing 5.1 μl of primer pool one or 5.6 μl primer pool two, 9.9 μl or 9.4 μl of template RNA, 25 μl of 2 x Platinum SuperFi RT-PCR Master Mix and 0.5 μl of SuperScript IV RT Mix. Thermocycling conditions consisted of 10 min at 55°C for reverse transcription, 2 min at 98°C to inactivate the reverse transcriptase enzyme followed by 40 cycles of 98°C for 10 s, 55°C for 10 s and 72°C for 3 min, with a final extension step at 72°C for 5 min. The sizes of mRT-PCR products were evaluated and amplicons were quantified on a TapeStation device (Agilent, CA, USA). Equal amounts of RT-PCR products from two reactions per sample were pooled together for downstream NGS library preparation and sequencing.

### DNAse treatment of viral RNA for multiplex RT-PCR

For samples with Ct values of 23 and higher, DNase treatment was also performed to remove human genomic DNA in the extracted viral RNA before doing multiplex RT-PCR. The DNA-free DNase treatment and removal reagents (Thermofisher, Australia) were used according to the manual provided by the manufacturer. Briefly, for 1 volume RNA, 1 μl DNase I and 0.1 volume of DNase I Buffer were added and incubated at 37°C for 30 min to digest DNA. Then, 0.1 volume of DNase Inactivation Reagent was added to inactivate DNase I at room temperature for 2 min followed by centrifugation to collect the supernatant for downstream analysis.

### Library preparation and sequencing on an Illumina platform

Two hundred ng of mRT-PCR products from each of two reactions per genome were pooled for the Illumina NGS library preparation. Pooled amplicons were cleaved into small fragments and tagged with adaptor sequences (also called on-bead tagmentation) followed by indexing according to manufacturer’s instruction. The pooled libraries were quantified by the Qubit dsDNA High Sensitivity (HS) assay (Thermo Fisher Scientific). The sizes of pooled libraries were measured with High Sensitivity D1000 ScreenTape and reagents (Agilent) on a TapeStation device (Agilent). Based on the results from the Qubit and TapeStation measurements, pooled libraries were diluted to 100 pM in Illumina RSB buffer, spiked with 1% PhiX, and sequenced on an Illumina iSeq sequencer using paired-end runs with 300 cycles (2×150 bp).

### Library preparation and sequencing on an Oxford Nanopore Technologies (ONT) platform

For nanopore sequencing, mRT-PCR products from each of two reactions per genome were purified by solid-phase reversible immobilization (SPRI) beads at a ratio of 0.5 (volume of beads /volume of PCR products) to remove small fragments. Three hundred ng of mRT-PCR products from each of two reactions per genome were pooled for the treatment with NEBNext Ultra II End repair and dA-tailing Module (New England Biolabs). According to the manufacturer’s protocols, the resulting products were firstly ligated with barcodes from the Native Barcoding Expansion kits by using NEB Blunt/TA Ligase Master Mix (New England Biolabs) and then with adapter from the Ligation Sequencing Kit (SQK-LSK109, Oxford Nanopore Technologies) added by using the NEBNext Quick Ligation Module (New England Biolabs). Finally, 50 fmol of library was loaded into a R9.4.1 FLO-MIN106 flow cell (Oxford Nanopore) and sequenced on a MinION Mk1B (Oxford Nanopore) for 3 hours.

### Illumina sequencing data analysis

An in-house bioinformatics pipeline was used for analyzing Illumina and ONT sequencing data, available at github [https://zenodo.org/record/6668599]. Briefly, the pipeline was written in the workflow management system snakemake ^28^. First, paired reads were processed with bbduk ^29^ for primer trimming using the following parameters: mink = 5, k = 12, hdist = 1. Second, the iterative refinement meta-assembler IRMA ^22^ was used for read trimming and filtering and reference-based assembly using a custom RSV module (available at github). Finally, quality control and error correction was performed by manually inspecting the assemblies and the alignment files.

### ONT sequencing data analysis (MinION-ONT)

The high-accuracy model was used for base-calling with MinKNOW software (Oxford Nanopore).

### Detecting human DNA reads in NGS datasets

We evaluated the abundance of human DNA reads in NGS datasets using Kraken 2 with the available Kraken Minidatabase containing the human genome and the full RefSeq set of bacterial and viral genomes ^23^.

### Phylogenetic Analysis

For whole-genome phylogenetic analysis, IQTree^30^ was run with automatic model selection (GTR+F+I+G4 model selected for both RSVA and RSVB) on samples with >95% coverage. In addition, ultrafast bootstrapping and SH-like approximate likelihood ratio test was performed (-bb 1000 -alrt 1000). Sequences were aligned to references (NC_001803.1 - RSVA, MG813995.1 - RSVB) with MAFFT (default settings)^31^. Finally, the phylogenetic trees were plotting using ggtree^32^. Additional sequences were retrieved from NCBI^33^. Clade references were retreived from Goya et al^34^.

### Statistical Analysis

GraphPad Prism software was used for statistical analyses. Comparisons between two groups were performed using the two-tailed *t*-test. *P* values less than 0.05 were considered as statistically significant and labelled as * in the figures. R software and packages were used for making graphs and curve fitting analyses.

### Data deposition

RSV genome sequences derived from viruses used in this study have been deposited into the GISAID database (https://gisaid.org/) with a full listing of viruses given in Supplementary table 2.

## Supplementary tables

Supplementary Table 1: Summary of NGS results for samples sequenced both on Illumina and ONT platforms

Supplementary Table 2: GISAID accession numbers and sample details

## Supplementary Figure

**Supplementary Figure 1.**
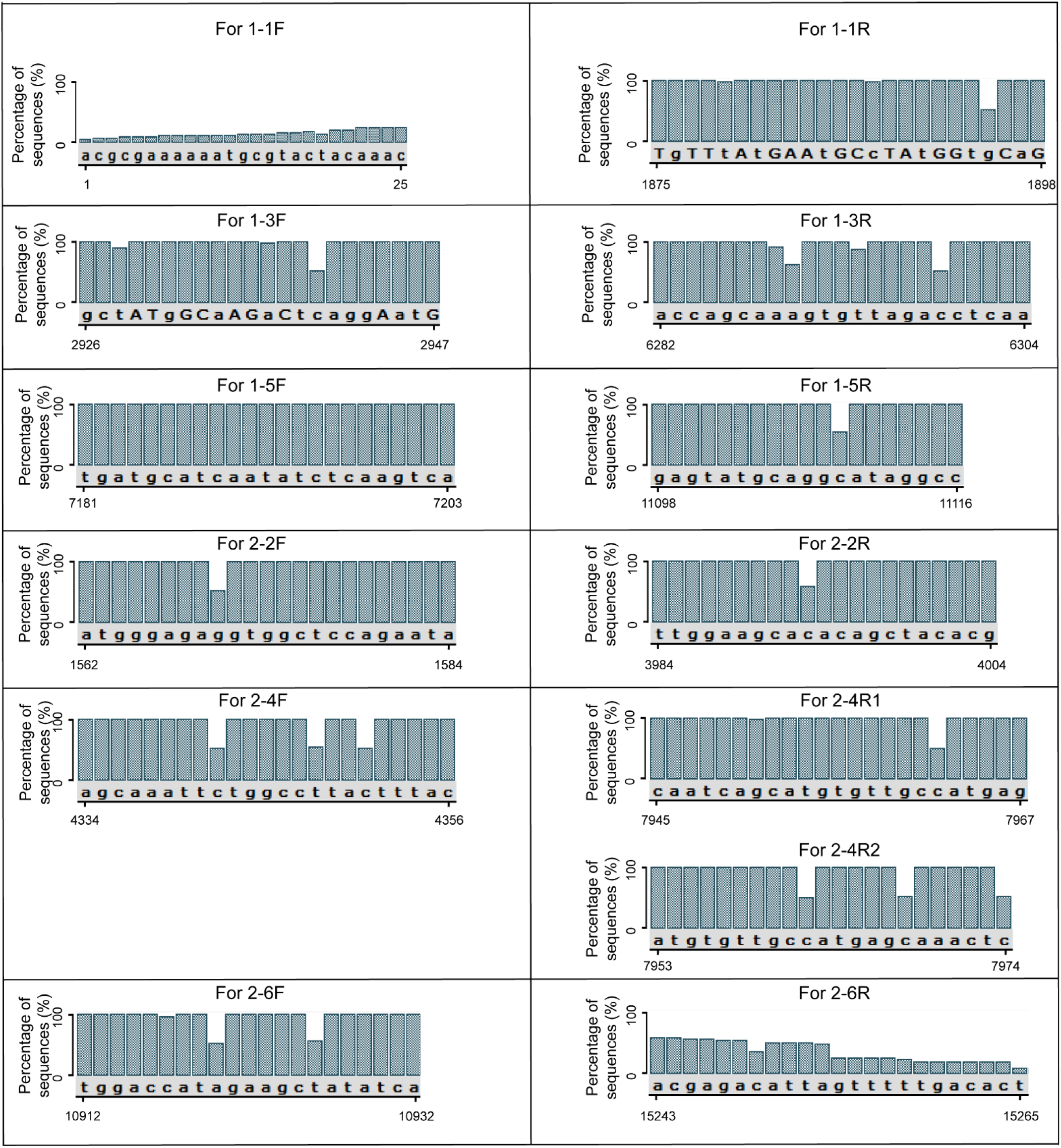
Alignment of RSV reference genomes for primer binding regions UGENE^35^ was used to align 1520 RSV-A and 1364 RSV-B genome sequences submitted to GISAID from 1^st^ January 2015 to 9th July 2021 with nucleotides more than 14900 bp. The location of primer binding regions in the consensus sequence generated by UGENE was based on a human RSV-B strain (GISAID accession numbers 2584506).

**Supplementary Figure 2.**
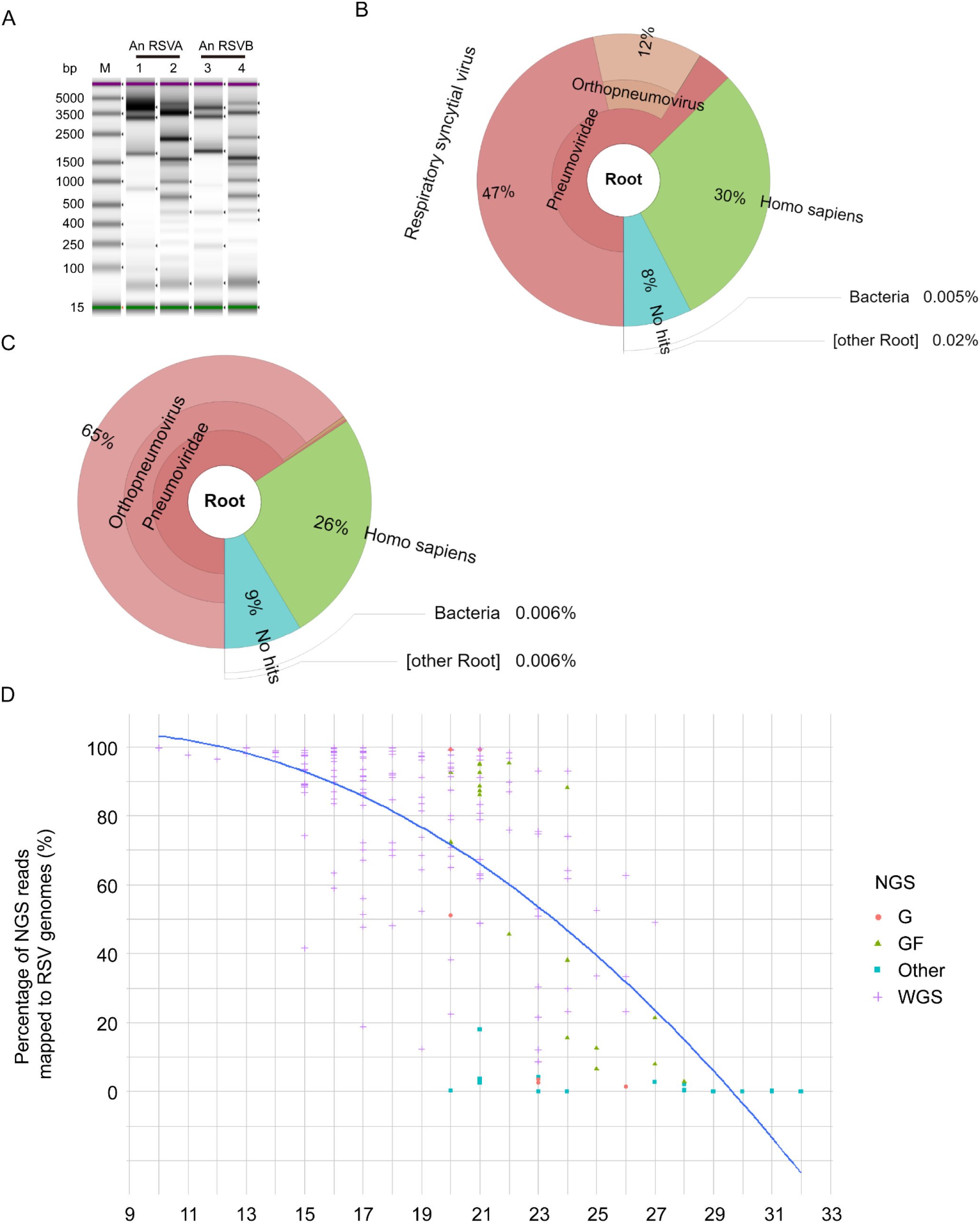
Samples directly tested with this new assay for sequencing RSV genomes (A) Tapestation images of two-pool PCR products of viral RNA from one RSV-A (hRSV/A/Australia/QLD-RBWH191/2021) and one RSV-B (hRSV/B/Australia/QLD-RBWH239/2021) positive clinical samples with unspecific amplicons observed. Read-level taxonomic classifications by Kraken for NGS results of RSV-A (B) and RSV-B (C) samples. (D) Dot plot showing the percentage of RSV specific sequencing reads against Ct values of samples. A blue line represents curve fitting to the percentage of mapped reads and Ct values. Coloured shapes represent different NGS results.

**Supplementary Figure 3.**
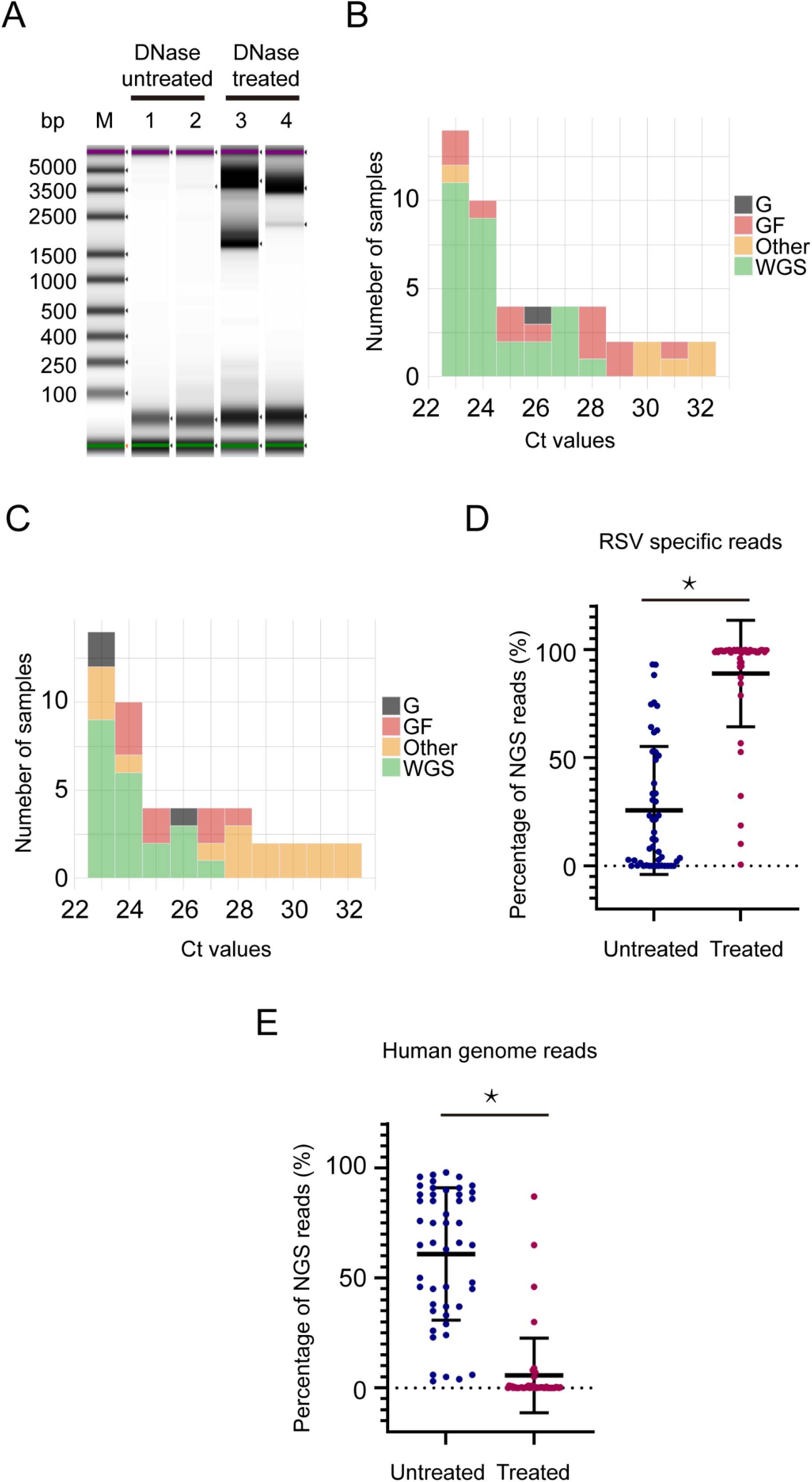
The impact of DNase treatment to viral RNA on the performance of this assay for sequencing RSV genomes (A) Tapestation images of RSV amplicons generated from one RSV-A positive clinical sample without and with DNase treatment. With (B) and without (C) human DNA removal, histograms of the Ct value distribution for clinical samples with RSV whole genome, *G* plus *F* genes, only *G* gene and missing *G* gene sequenced (also called “other”), respectively. (D) Dot plot showing the percentage of RSV specific reads from sequenced samples without and with human genome background removal by DNase treatment. (E) Dot plot showing the percentage of human reads from sequenced samples without and with human genome background removal by DNase treatment. Data are expressed as mean ± SD. *t*-test analysis was performed for statistical significance. *P* values less than 0.05 were considered as statistically significant and labelled as * in the figures.

**Supplementary Figure 4.**
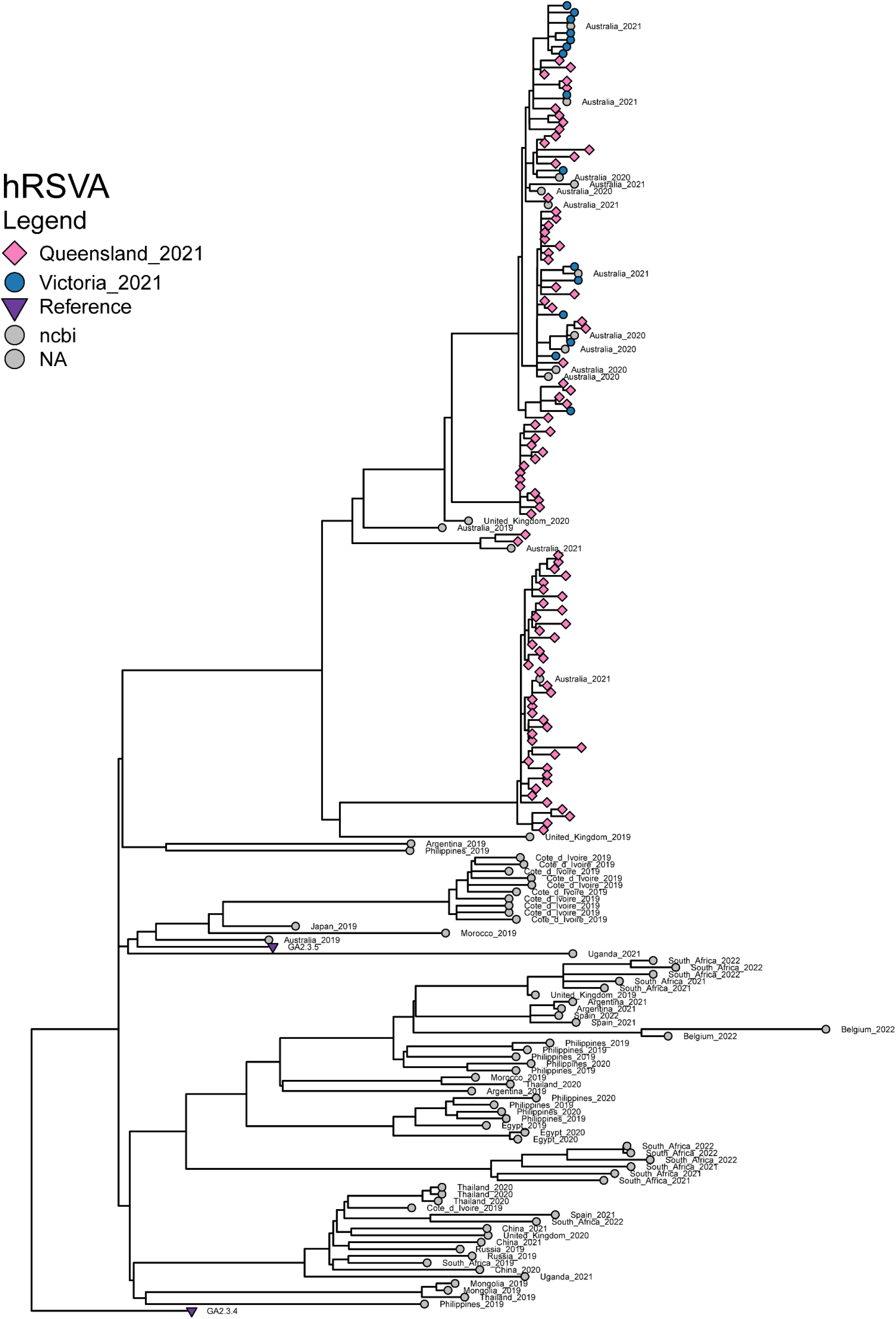
Phylogenetic analysis of WGS of RSV-A generated using the 2-tube method described in this method.

**Supplementary Figure 5.**
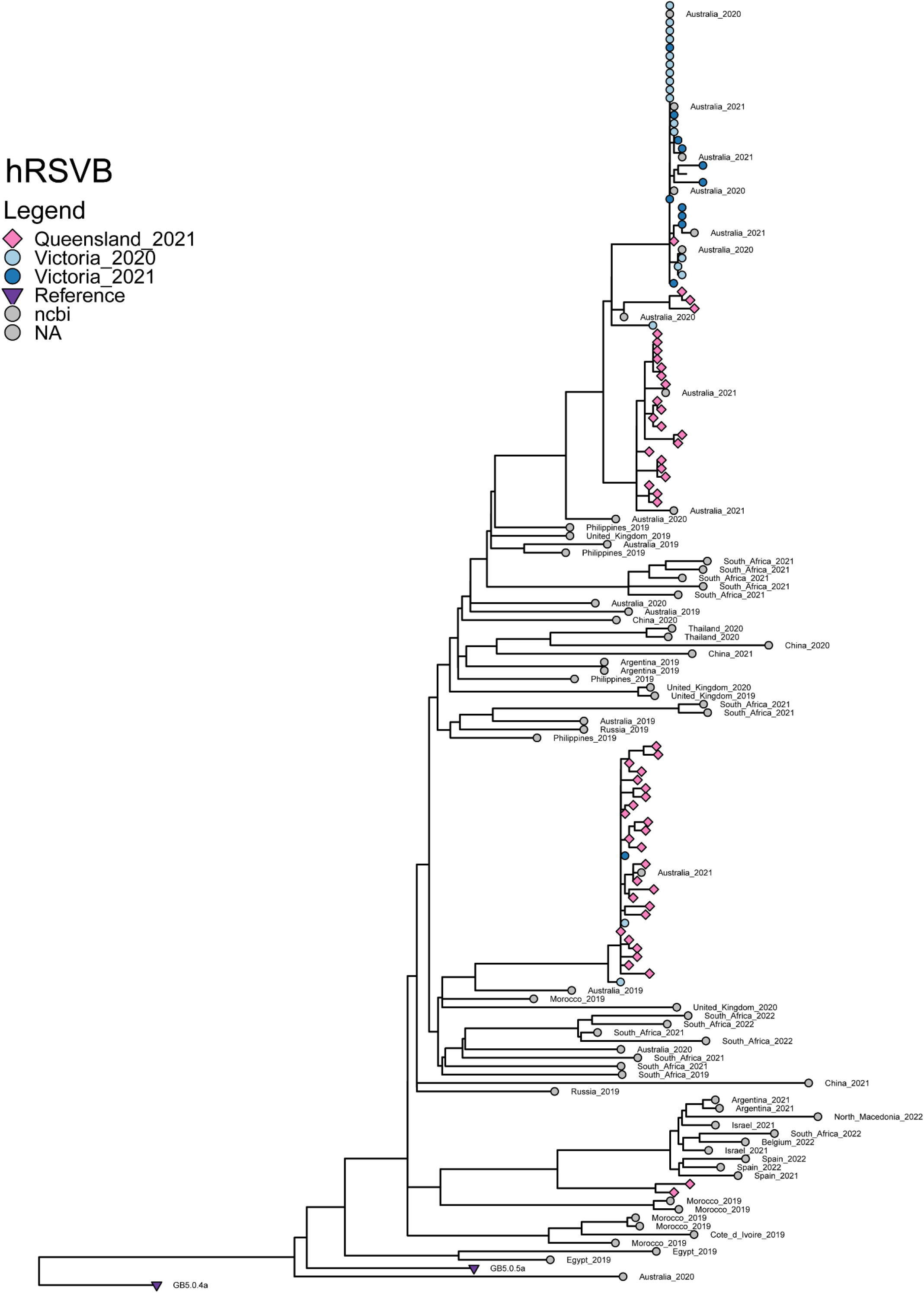
Phylogenetic analysis of WGS of RSV-B generated using the 2-tube method described in this method.

## Acknowledgments

The WHO Collaborating Centre for Reference and Research on Influenza is supported by the Australian Government Department of Health.

